# The TUDOR domain of SMN is an H3K79^me1^ histone mark reader

**DOI:** 10.1101/2022.10.06.511070

**Authors:** Olivier Binda, Aimé Boris Kimenyi Ishimwe, Maxime Galloy, Karine Jacquet, Armelle Corpet, Amélie Fradet-Turcotte, Jocelyn Côté, Patrick Lomonte

## Abstract

Spinal Muscle Atrophy (SMA) is the leading genetic cause of infant mortality and results from the loss of functional Survival Motor Neuron (SMN) protein by either deletion or mutation of the *SMN1* gene. SMN is characterized by a central TUDOR domain, which mediates the association of SMN with arginine methylated (R^me^) partners, such as COILIN, FIBRILLARIN, and RNApolII. Herein, we biochemically demonstrate that SMN also associates with histone H3 monomethylated on lysine 79 (H3K79^me1^), defining SMN as the first known H3K79^me1^ histone mark reader, and thus the first histone mark reader to recognize both methylated arginine and lysine residues. Mutational analyzes provide evidence that SMN_TUDOR_ associates with H3 via an aromatic cage. Importantly, most SMN_TUDOR_ mutants found in SMA (SMN_ST_) patients fail to associate with H3K79^me1^.

**Summary Blurb:** Spinal Muscle Atrophy (SMA) is caused by mutation or deletion of *SMN1* gene. Survival Motor Neuron (SMN) protein associates with histone H3 mono-methylated on lysine 79 (H3K79^me1^) through its central TUDOR domain. SMA-linked mutations occur within the TUDOR domain and prevent association with histone H3.

## Introduction

Although the loss of *SMN1* was found in 1995 as responsible for the monogenic pathology SMA (1), it took until 2019 to develop a gene therapy treatment approved by the FDA (2). However, long-term efficacy of such treatment remains unknown and not appropriate to all patients [reviewed (3,4)]. It is thus critical to continue basic research on SMN biochemistry for SMA patients and for general knowledge, which is likely to impact on other neurological disorders with common biochemical pathways, such as amyotrophic lateral sclerosis (ALS) and Fragile X syndrome.

SMN, the protein, also known as GEMIN1, is principally recognized as a marker of membrane-less nuclear structures called Cajal bodies, first identified in neuronal tissues by Santiago Ramón y Cajal (5) and recently found to phase separate (6). SMN most documented cellular function is to assemble snRNPs and thus regulate RNA metabolism and splicing [reviewed in (7,8)]. Essentially, SMN cellular functions are centered on its TUDOR domain, which coordinates protein-protein interactions with arginine methylated (R^me^) proteins, such as COILIN (9,10), RNApolII (11), and other R^me^GG motif proteins (6,12,13).

In human there are 2 copies of the gene that encodes SMN, namely *SMN1* and *SMN2*. The *SMN1* gene is lost in most SMA patients, leading to low levels of SMN protein, but approximately 10% of SMA cases harbor mutations. Interestingly, most SMN mutations congregate either within the carboxy terminal oligomerization domain or within the central TUDOR domain [reviewed (4)], suggesting an important role for the TUDOR domain and the capacity of SMN to oligomerize in the maintenance of motor neuron homeostasis. The TUDOR domain is part of a large family of histone and non-histone recognizing modules that associate with post-translationally modified or unmodified proteins (14,15). Previous work from our team suggests that SMN associates with histone H3 and localizes with damaged centromeres in a DOT1L methyltransferase-dependent manner (16). Although DOT1L is often depicted as the only histone H3 lysine 79 (H3K79) methyltransferase (17,18), *Dot1l*^-/-^ knockout cells retain H3K79^me2^ albeit at extremely low levels (0.5% in *Dot1l*^-/-^ versus 3.3% H3K79^me2^ in wild-type cells) (19), suggesting that there may be other methyltransferase(s) capable of modifying H3K79. There are indeed a few studies suggesting that NSD1 and NSD2 methyltransferases could mono- and di-methylate H3K79 (20,21).

Histones and histone post-translational modifications (histone marks) are central to chromatin signalling pathways. Essentially, genomic DNA is wrapped around small basic proteins called histones to form nucleosomes, a repetitive unit constituting the chromatin framework, which regulates access to genetic information. Generally, histone modifications such as H3K4^me3^, H3K9^me1^, and H3K79^me1^ mark the chromatin for access to the genetic information, while modifications such as H3K9^me3^ and H3K27^me3^ marks restrict access to the genetic information (22). Regarding H3K79^me1^, the mark correlates with alternative splicing paterns between cell lines (23), in agreement with the positioning of H3K79^me1^-marked nucleosomes on exons (24). More precisely, the H3K79^me1^ and H3K79^me2^ marks are found at alternative 3’
s and 5’ splice sites (25). The H3K79^me1^ mark is the most prominent state of methylation on H3K79 in mouse ES cells (19). Notably, the mouse model of *Dot1l*^-/-^ is embryonic lethal while *Dot1l*^-/-^-derived cells harbor alternative lengthening of telomere (ALT) phenotype (19). Moreover, DOT1L plays an important role in neuronal development (26).

Herein, using biochemical approaches, purified recombinant proteins, and recombinant nucleosomes, we define SMN_TUDOR_ as the first H3K79^me1^ mark reader, and also the first histone mark reader capable of reading both R^me^ and K^me^ states. Importantly, SMA-linked SMN mutations (SMN_ST_) prevent SMN-H3 interactions, suggesting the involvement of chromatin signalling pathways in SMA genetic pathology.

## Results

### Purified recombinant SMN interacts directly with histone H3 in vitro

We previously reported that SMN associates with damaged centromeres in a DOT1L-dependent fashion (16), suggesting that SMN could associate with histone H3 methylated on lysine 79. Indeed, initial investigations suggested that SMN associates with H3K79^me2^ peptides (16). Herein, we thus aimed to further characterize biochemically this potential interaction. Using recombinant human SMN purified from *Escherichia coli* by a GST affinity purification scheme, GST-SMN was subjected to pulldown assays in the presence of calf thymus histones (CTH), a classic source of modified histones (27). In agreement with previous work (16), GST-SMN was capable to associate directly with histone H3 while the GST alone control failed to associate with histones (**Figure 1A**). Similar experiments were conducted with the PHD domain of the H3K4^me3^ reader ING3 (ING3_PHD_) and a characterized aromatic cage mutant known to be unable to associate with H3 (ING3_W385A_) (28,29), as positive and negative controls, respectively. These pulldowns were analyzed by immunoblotting against core histones and histone variants. As expected, ING3_PHD_ associated with H3, while the aromatic cage mutant ING3_W385A_ failed to do so (**Figure 1B**). As seen in Figure 1A, we observed that GST-SMN associates predominantly with H3 (**Figure 1B**). Extended immunoblots for each pulled down proteins are provided in supplementary materials (**Figure S1A**). These experiments validate the association of SMN with histone H3.

**Figure 1:**
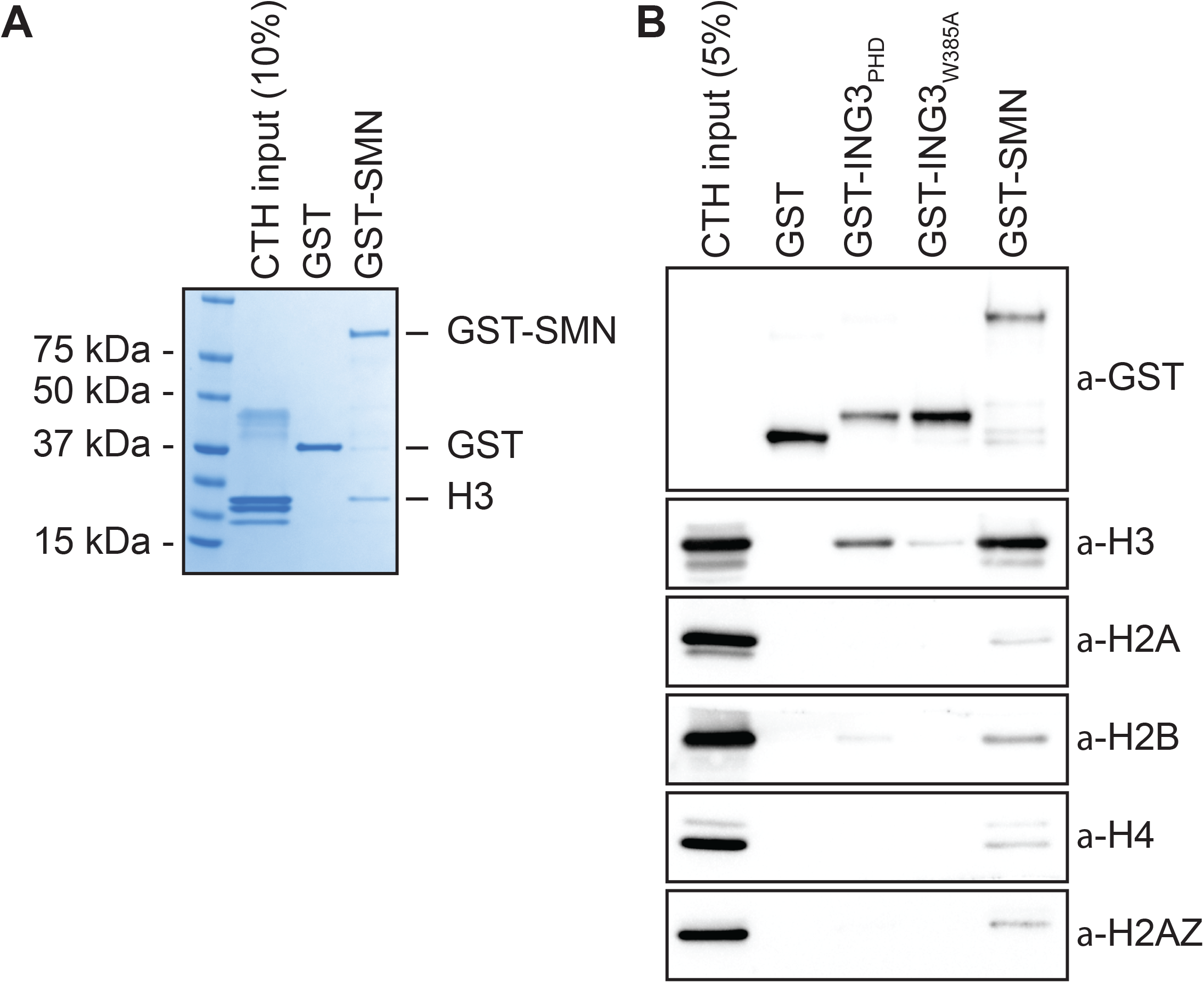
SMN associates with histone H3. (**A**) GST alone or GST-tagged SMN were used in GST-pulldown assays in the presence of calf thymus histones (CTH). Pulldowns were analyzed by SDS-PAGE followed by Coomassie staining. (**B**) As in panel A, but pulldowns were analyzed by immunoblotting using the indicated antibodies.

### An intact TUDOR domain is required for SMN to interact with H3

As SMN harbors a TUDOR domain, which is found in several other histone mark readers (30), we then set out to define the minimal region of SMN required for H3-binding and generated a panel of truncated forms (**Figure 2A**). Using these, we found that deletion of either the amino terminus (SMN_ΔN_) or the carboxy terminus (SMN_ΔC_) affected the association with H3 minimally (**Figure 2B**). However, the TUDOR domain on its own (SMN_TUDOR_) failed to associate with H3 as well as the full-length form of SMN, but seemed to be required for the interaction since the amino terminus (SMN_Nterm_) and carboxy terminus (SMN_Cterm_), which lack the TUDOR domain, only bound weakly to H3 (**Figure 2B**). Thus, we conclude that the TUDOR domain is required *in vitro*, but not sufficient for SMN to associate with H3. We then extended the TUDOR domain on both sides, with actual SMN wild-type sequences, and found that an extension by 25 amino acid residues on its amino terminal side restored to some extend the association of SMN with H3 (**Figure 2C**). Extension on the carboxy terminal end of SMN_TUDOR_ by 5, 10, or 15 residues did not appear to improve the association of SMN to histone H3 (**Figure S1B**). We conclude that SMN_TUDOR_ is necessary and sufficient for association with histone H3, but requires additional residues outside the classical defined TUDOR domain.

**Figure 2:**
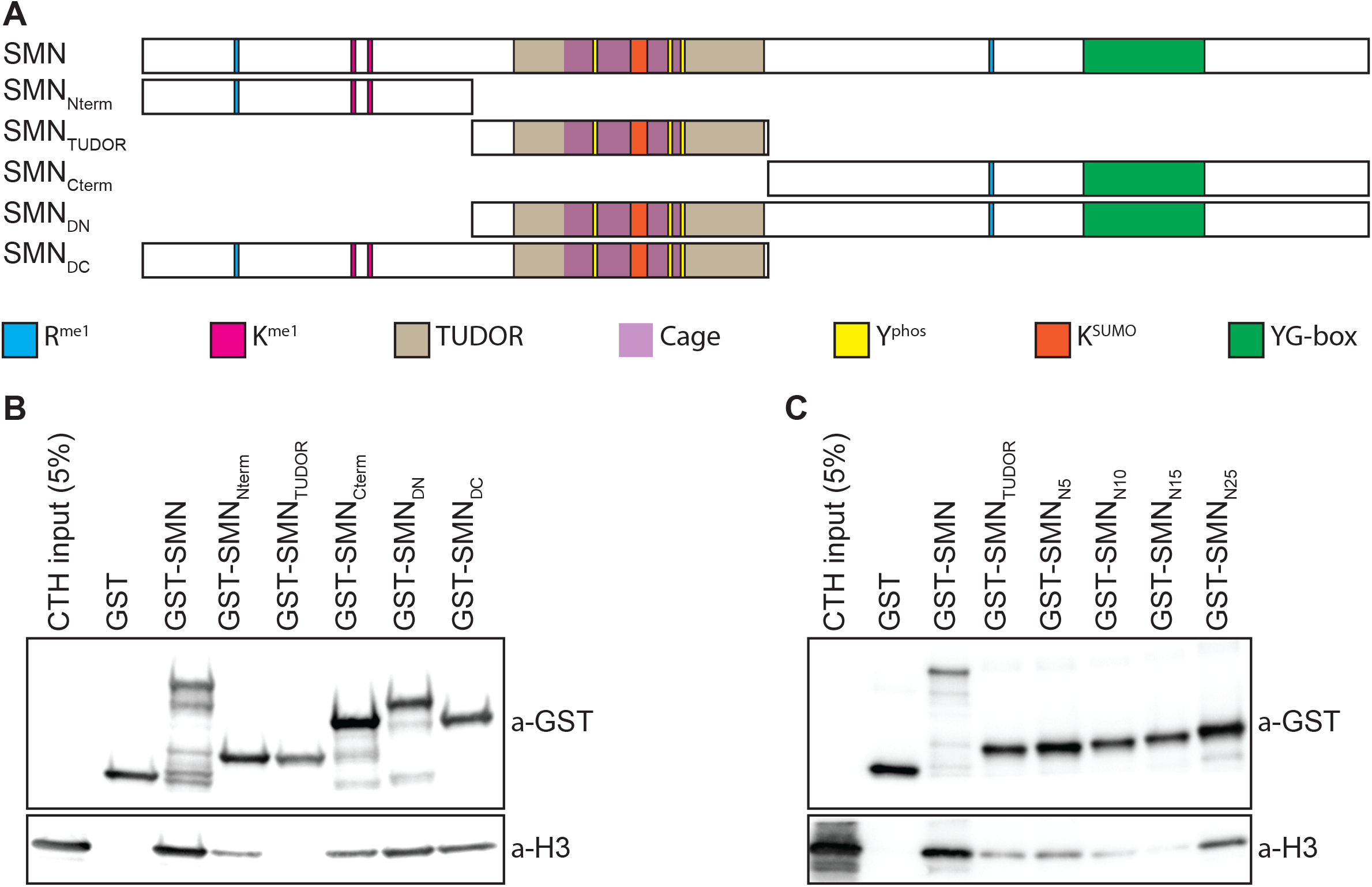
The TUDOR domain is required but not sufficient for SMN to associate with H3. (**A**) Schematic representation of SMN truncated forms used to assess the region of SMN responsible for the association with H3. Post-translational modification (methyl-arginine [R^me^], methyl-lysine [K^me^], phospho-tyrosine [Y^phos^], and SUMOylated lysine [K^SUMO^]) sites are highlighted. The aromatic cage within the TUDOR is marked in purple and the dimerization YG-box highlighted in green at the carboxy terminus. (**B**) GST-pulldowns were performed as in Figure 1 and analyzed by immunoblotting with α-GST and α-H3 antibodies. (**C**) As in panel B, but with amino terminal extensions on recombinant SMN_TUDOR_.

The solution structure of SMN bound to R^me^ residue displays an aromatic cage composed of W102, Y109, Y127, and Y130 (31). Aromatic cages are broadly found in histone mark readers and involved in sensing methylation states (30). To demonstrate the importance of the TUDOR domain in mediating the interaction between SMN and H3, we have mutated W102, Y109, Y127, and Y130 aromatic cage sites (SMN_AC_ mutants) individually to alanines and performed pulldown assays to assess the interaction of SMN mutants with H3. Unexpectedly, unlike other readers such as ING4 (32), HP1α (33), or MPP8 (33), single mutations to alanine within the aromatic cage did not appear to alter SMN_AC_ binding to H3 and retained the capacity to associate with H3 (**Figure 3A**). We thus mutated these four residues in various combinations and found that W102 and Y130 seemed to be the most important residues of the aromatic cage as all the SMN forms that had reduced binding affinity to H3 contained W102A and Y130A mutations (**Figure 3B-C**). Together, these results demonstrate that SMN associates directly with H3 and requires an intact TUDOR domain aromatic cage.

**Figure 3:**
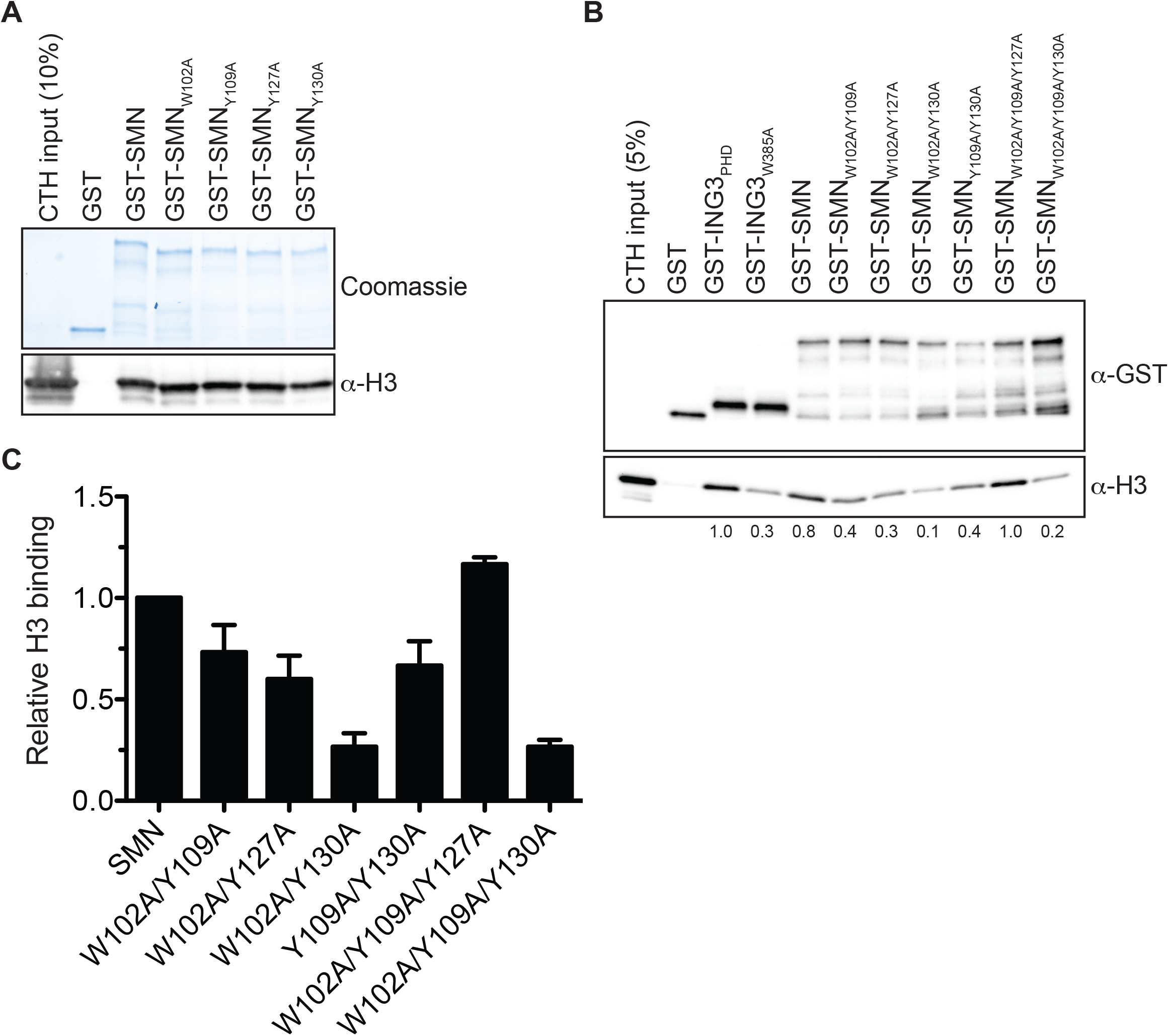
The aromatic cage within SMN_TUDOR_ is critical for SMN-H3. (**A**) GST-pulldowns were performed as in Figure 1. (**B**) As in panel A, but GST-ING3_PHD_ and ING3_W385A_ were used as positive and negative controls, respectively. (**C**) The level of H3 was measured using Image Lab (Bio-Rad Laboratories) from 3 independent experiments.

### SMA-linked SMN_TUDOR_ mutants fail to associate with H3

In about 10% of SMA cases, *SMN1* is not deleted, but mutated. These mutations aggregate mostly in the dimerization domain or within the TUDOR domain [reviewed in (4)]. Given that SMN associates with H3 through an aromatic cage within its TUDOR domain (**Figures 2-3**), we investigated whether SMA-linked TUDOR mutations (SMN_ST_) impact on the association of SMN with histone H3. Experiments with the aromatic cage mutant SMN_Y109C_ and the other SMA-linked mutant SMN_E134K_ seemed to show that E134K minimally impact the capacity of SMN to interact with H3 (**Figure 4A**). We thus expanded our panel to include all known SMN_ST_ (4). Interestingly, SMN_ST_ W92S, G95R, A111G, and I116F had impaired capacity to associate with histone H3, while V94G, Y109C, Y130C, E134K, and Q136E retained approximately wild-type level of binding to H3 (**Figure 4B**). These results suggest that SMA may arise in some cases from impaired SMN-H3 interactions or other interactions requiring an intact SMN_TUDOR_.

**Figure 4:**
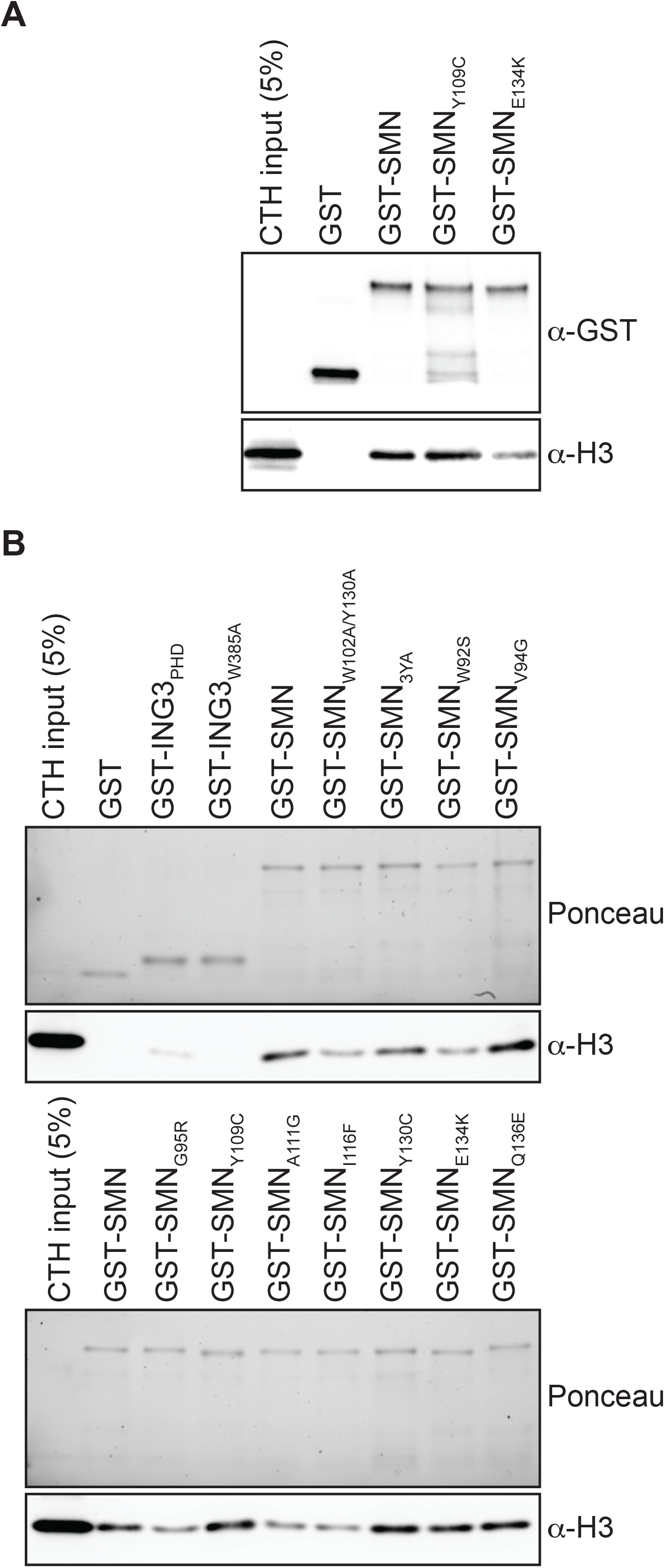
SMA-linked SMN_TUDOR_ mutants impact SMN interaction with H3. (**A**) GST-pulldowns were performed with full-length recombinant GST-SMN and SMA-linked mutants GST-SMN_Y109C_ and GST-SMN_E134K_. (**B**) As in panel A, but with a complete panel of TUDOR mutants. The pulldowns were analyzed by immunoblotting against histone H3.

### Defining SMN as the first H3K79^me1^ reader

Previous work suggests that SMN may associates with H3 sequences surrounding the lysine 79 methylation site (16). We thus analyzed GST-SMN pulldowns by immunoblotting against methylated H3K79 forms and found that the H3 species that associate with SMN are predominantly the H3K79^me1^ and H3K79^me2^ forms (**Figure 5A**). Although controversial, another TUDOR domain protein, 53BP1, was also reported to associate with methylated H3K79 (34,35). We thus investigated how SMN associates to histones compared to 53BP1 and found that SMN bound to H3 with the H3K79^me2^ mark and modestly to H4 with the H4K20^me2^ modification at least as well as 53BP1 if not better (**Figure 5B**). We then used the H3K4^me3^ reader ING3_PHD_ as a control to assess the enrichment of the H3K79 methyl marks by SMN. We found that the H3 pulled down by SMN was enriched with the H3K79^me1^ mark compared to ING3_PHD_ (**Figure 5C**), suggesting that SMN may associate with this mark (i.e. H3K79^me1^). Hitherto, our results show that SMN associates with H3 harbouring the H3K79^me1/me2^ marks, but does not demonstrate that SMN associates with the H3K79^me1/me2^ marks themself.

**Figure 5:**
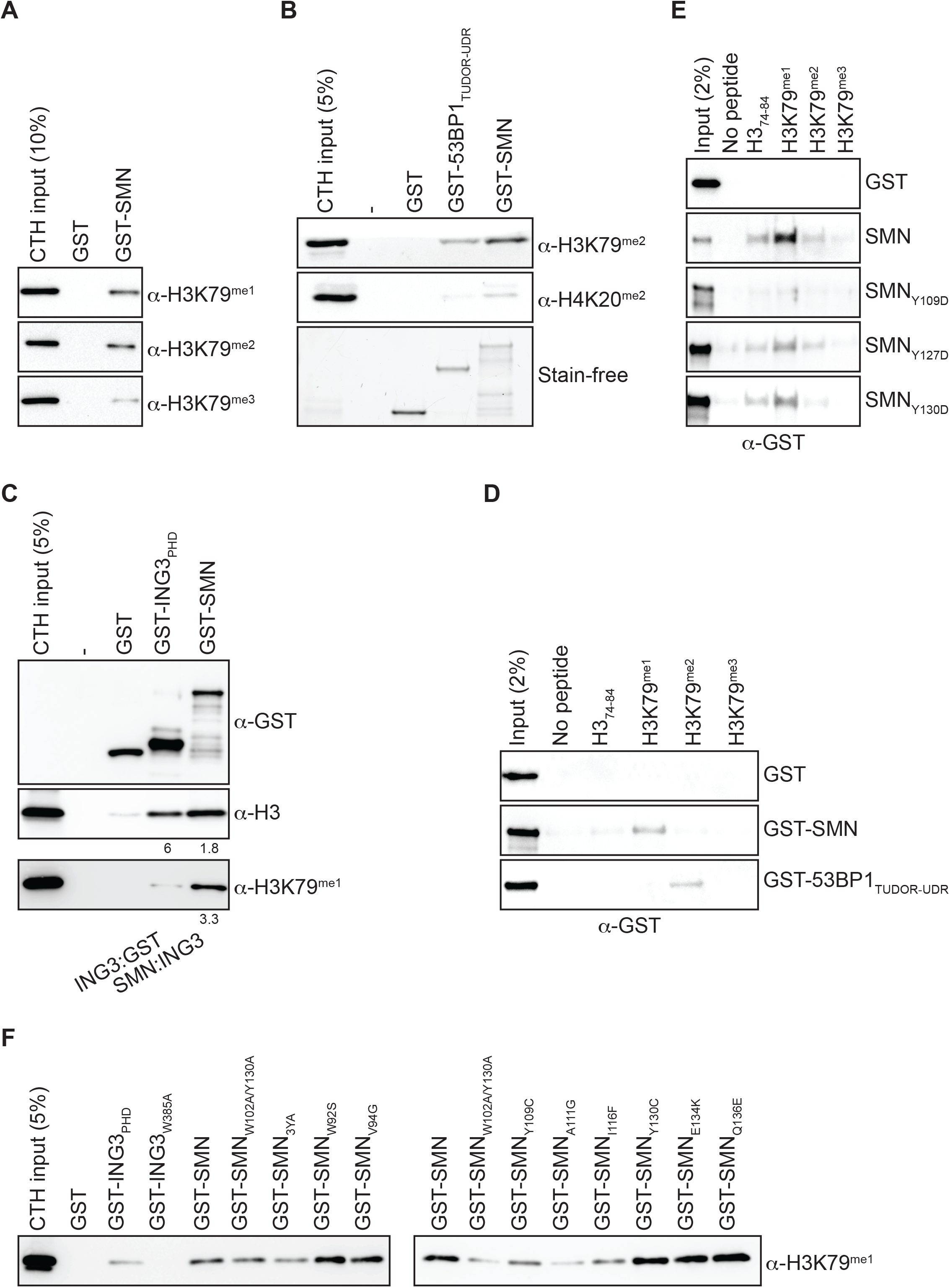
SMN is a H3K79^me1^ reader. (**A**) GST-pulldowns with CTH were analyzed immunoblotting using α-H3K79^me1^, H3K79^me2^, or H3K79^me3^ antibodies. (**B**) As in panel A, but GST-53BP1_TUDOR-UDR_ was used as a positive control known to associate with H3K79^me2^ (34) and H4K20^me2^ (35). (**C**) As in panel A, but GST-ING3_PHD_, a H3K4^me3^ reader, was used as a negative control. Ratios of ING3:GST and SMN:ING3 signals are indicated under the immunoblots. (**D**) Biotinylated synthetic histone peptides were pulled down using streptavidin-sepharose in the presence of GST, GST-SMN, or GST-53BP1_TUDOR-UDR_. Pulldowns were analyzed by immunoblotting using an α-GST antibody. (**E**) As in panel D, but biotin H3K79 peptides were pulled down in the presence of GST-SMN aromatic cage mutants. (**F**) As in Figure 4, panel B, but the same samples were analyzed with an α-H3K79^me1^ antibody.

As mentioned above, although SMN associates with H3 methylated on lysine 79 (**Figures 4** and **5**), our results do not demonstrate that SMN associates directly with the H3K79 methylated marks. We thus performed pulldowns using synthetic biotinylated peptides, which are either unmodified, mono-, di-, or trimethylated on H3K79 and recombinant SMN. While performing various contraptions using peptide pulldown assays, we found that SMN associates preferentially with H3K79^me1^, while 53BP1_TUDOR_ bound to the H3K79^me2^ peptide (**Figure 5D**). We thus conclude so far that SMN associates directly with H3K79^me1^, at least *in vitro* using biochemical assays.

Interestingly, the tyrosine residues (Y109, Y127, and Y130) of the aromatic cage are reported to be phosphorylated (36). Tyrosine phosphorylation within aromatic cages is hypothesized to regulate reader-mark interactions (33). Conversion of Y109, Y127, or Y130 to aspartic acid, to mimic the negative charge of the phosphate moiety, appears to reduce to binding of SMN to H3K79^me1^ (**Figure 5E**), suggesting that Y^phos^ of SMN_TUDOR_ could regulate the association of SMN with methylated partners, such as H3. Our *in vitro* biochemical characterization identifies SMN as the first known H3K79^me1^ histone mark reader.

As we defined SMN as an H3K79^me1^ reader, we thus then assessed the impact of SMN_ST_ on binding to H3 methylated on K79. Similarly to H3 (**Figure 4B**), W92S, Y109C, A111G, and I116F had an impact on the association of SMN with H3K79^me1^, while V94G, G95R, Y130C, E134K, and Q136E had no apparent effect on the binding of SMN (**Figure 5F**).

The H3K79^me1^ mark is, unlike histone modifications, such as H3K4^me3^ and H3K9^me3^, not found on the unstructured histone tail, but within the histone H3 core region (37). Thus, the nucleosome inherent structure may impact on the accessibility of H3K79^me1^ to potential readers, such as SMN. To assess this possibility, we generated recombinant nuclear core particles (rNCP) containing histone H3 containing mutations C110A and K79C (K_C_79) to specifically modify K_C_79 with a monomethyl-nucleoside analog. Unmodified H3K_C_79 (rNCP-H3K_C_79^me0^) or monomethylated H3K_C_79 (rNCP-H3K_C_79^me1^) were used to reconstitute octamer and subsequently nucleosome assembly (**Figure S2**). These were used in GST-SMN pulldown experiments, which confirmed that SMN associates with H3K79^me1^ within the nucleosomal context (**Figure 6**). Specifically, SMN associated with rNCP-H3K_C_79^me1^, but not with unmodified rNCP-H3K_C_79^me0^ form (**Figure 6**). Moreover, most SMA-linked SMN_TUDOR_ mutants failed to associate with H3K_C_79^me1^ nucleosomes (**Figure 6**). Specifically, W92S, G95R, Y109C, A111G, I116F, and Y130C mutants prevent SMN from associating with H3K_C_79^me1^ nucleosomes, while V94G, E134K, and Q136E had no discernable impact on the SMN-H3K79^me1^ interaction (**Figure 5D**). Interestingly, the association of SMN and SMN_ST_ with free H3, peptides, and rNCP varied slightly (**Table S1**). More precisely, G95R bond H3K79^me1^-marked H3, but not the rNCP-H3K_C_79^me1^, while Y109C and Y130C bound total H3 from calf thymus, but not rNCP-H3K_C_79^me1^.

**Figure 6:**
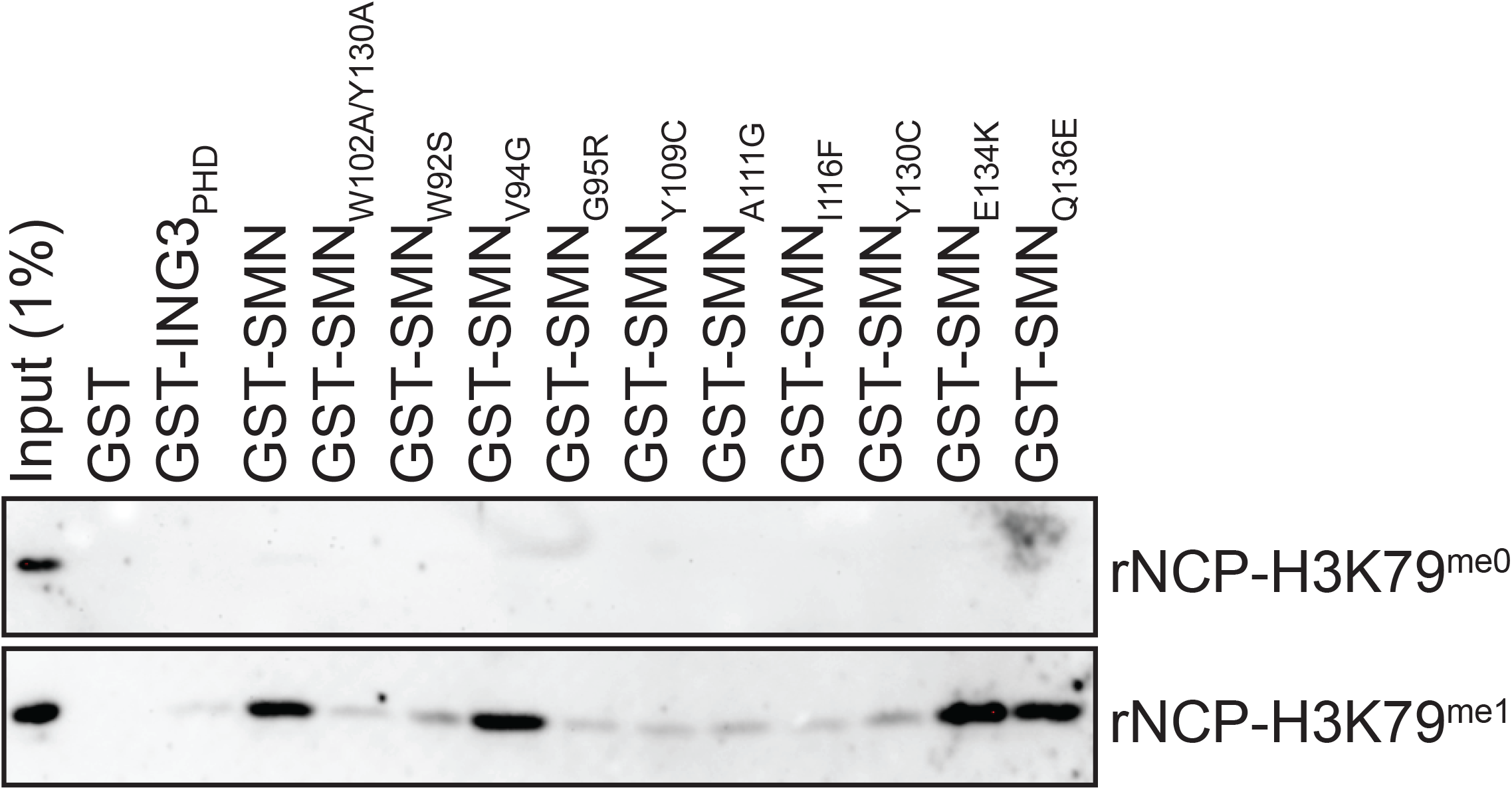
SMN associates with H3K_C_79^me1^ nucleosomes. Recombinant nucleosomes core particles were assembled and chemically modified on lysine 79 (rNCP-H3K_C_79^me0^ and rNCP-H3K_C_79^me1^). Indicated GST-tagged proteins were pulled down using glutathione-sepharose in the presence of either rNCP and analyzed by immunoblotting using an α-H3 antibody.

## Discussion

Unlike histone tail modified residues, such as H3K4, H3K9, or H3K27, histone H3 lysine 79 is found within the core of histone H3 with the side chain of K79 sticking out like a broken bicycle wheel spoke. Specifically, methylation of H3K79 (H3K79^me2^) alters the surface of the nucleosome (38).

Although H3K79^me2^ can be weakly recognized by the tandem TUDOR domain (TTD) of 53BP1 (34), 53BP1_TTD_ prefers H4K20^me2^ with a K_D_ of 20 μM compared to 2 mM for H3K79^me2^ (35). Another study found that H3K79^me3^ can be recognized *in vitro* by the WD40 domain of EED with low affinity (K_D_ of > 400 μM) [PDB 3JZH (39)]. Although the TUDOR domain protein Fragile X Mental Retardation Protein (FMRP) associates with H3K79^me2/me3^ with a K_D_ of 135 nM, it also binds to H3K4^me2/me3^, H3K9^me3^, H3K27^me1/me2/me3^, and H3K36^me2/me3^ (40). However, the FMRP family members FXR1 and FXR2 fail to associate with H3K79^me3^, but rather have affinity for H4K20^me3^ (41). Finally, there is also HDGF2_PWWP_ that can recognize H3K79^me3^ (42). Regardless of all these readers that can recognize H3K79^me2/me3^, no reader for H3K79^me1^ have been identified so far. Previous work from our laboratory described a DOT1L-dependent relocation of SMN in response to damaged centromeres that required an intact TUDOR domain (16), suggesting that SMN_TUDOR_ may interact with methylated H3K79. We herein define using multiple assays and approaches that SMN_TUDOR_ associates with H3K79^me1^.

Interestingly, SMA-linked SMN_TUDOR_ mutants (SMN_ST_) impact differently the interaction of SMN with free histone H3, K79^me1^-modified H3, and H3K_C_79^me1^ rNCP (summarized in **Table S1**). We can only speculate on the nature of the discrepancies between the different histone contexts (i.e. peptides, free histones, rNCP), but likely oligomerization and/or nucleotide-binding properties of SMN play a role.

Although several readers recognize dual marks, such as ZMYND8_PHD/BD_-H3K4^me1^K14^ac^ (43) and SETDB1-H3K9^me3^K14^ac^ (44), while others have affinity for a handful of sites, such as PHD that recognize unmodified histone H3, H3K4^me^, or H3K9^me^ (45,46), few readers recognize both methylated arginine and lysine residues indiscriminately. For instance, SPIN1 recognizes H3R8^me2a^ with TUDOR-like SPIN repeat 1 and H3K4^me3^ via SPIN repeat 2 (47), but also H3K9^me3^ and H3K4^me3^ (48). Interestingly, the overall structures of H3K4^me3^R8^me2a^-or H3K4^me3^K9^me3^-bound SPIN1 remain similar, except for a small rotation of W72 and F251 side chains in the aromatic cage of SPIN repeat 1 to accommodate either H3K9^me3^ or H3R8^me2a^ (48). We conclude that SMN_TUDOR_ likely conforms like SPIN1 to be able to associate with H3K79^me1^ as well as R^me^GG-containing proteins (e.g. COIL, FBL).

## Conclusions

SMN_TUDOR_ is herein identified as the first reader of the H3K79^me1^ mark, which is found on exons and linked to alternative splicing events. Importantly, SMA-linked SMN_TUDOR_ mutants (SMN_ST_) impact profoundly interactions with histones. Since the TUDOR royal family has always been promiscuous, it is interesting to discover that SMN_TUDOR_ is bounding with both methylated arginines and methylated lysines.

## Methods

### Recombinant protein expression

The cDNA of human full-length SMN and truncations were inserted in pGEX-6P-1 (GE Healthcare) using *BamHI* and *XhoI*. Aromatic cage mutants were generated by site-directed mutagenesis using Pfu Turbo (Stratagene) followed by *DpnI* (NEB) digestion. Constructs were sequence-verified (GATC Biotech AG or Biofidal) and transformed into BL21 DE3 cells (Stratagene). BL21 cells were grown overnight with ampicillin selection at 37°C with agitation. The following day, cultures were scaled up in 250 mL LB (Sigma) and grown at 37°C until OD_600_ ∼0.6. Then, expression of recombinant GST proteins was induced with 0.2 mM IPTG for 2.5-3 hours at 37°C. Cells were harvested by centrifugation, resuspended in lysis buffer (50 mM Tris pH 7.5, 150 mM NaCl, 0.05% NP-40, supplemented Complete EDTA-free [Roche]). After a brief sonication, lysates were cleared by centrifugation, and incubated with glutathione-sepharose (GE Healthcare) at 4°C on a tumbler wheel. After extensive washing, GST proteins were eluted with 10 mM reduced glutathione (Sigma) in 50 mM Tris pH 8.0.

### Antibodies

The α-H3 (ab1791), α-H4K20me2 (ab9052), and HRP-conjugated α-GST (ab3416) antibodies were obtained from Abcam. The α-H2A (07-146), α-H2B (07-371), and α-H4 (62-141-13) were obtained from Millipore. Methyl-specific H3K79^me1^ (pAb-082), H3K79^me2^ (pAb-051), and H3K79^me3^ (pAb-068) antibodies were purchased from Diagenode. The α-H2AZ antibody was described elsewhere (49).

### Histone interactions

GST pulldowns were performed with 25 μg calf thymus histones (Worthington) and ∼1 μg recombinant GST or GST-SMN in freshly made 25 mM bis tris propane (Sigma B6755) pH 6.8 buffer with 1 M NaCl and 0.05% NP-40. Glutathione-sepharose beads (GE17-5132-01) were added for an hour, then washed 4 times with 1 mL bis tris propane buffer.

### Peptide pulldowns

Peptide pulldowns were performed with 1 μg biotinylated H3 peptides (16) and ∼1 μg recombinant GST or GST-SMN in freshly prepared 25 mM bis tris propane buffer with 200 mM NaCl and 0.05% NP-40. Streptavidin-sepharose beads (GE17-5113-01) were added for an hour, then washed 4 times with 1 mL bis tris propane buffer.

### H3K79 recombinant nucleosomes

Bacterial expression vectors for histones H2A and H2B were purchased from Addgene (42634 and 42630, respectively). Plasmids to express *X. laevis* H3 (xH3) in pET-3d and xH4 in pET-3a were obtained from Professor Arrowsmith (University of Toronto, Canada). Introduction of C110A and K79C mutations in xH3 were done by site-directed mutagenesis using QuikChange (Stratagene) and the plasmid was sequence-verified. Recombinant histones were purified from *E. coli*, and modified where indicated, prior to octamer assembly and subsequent refolding of recombinant nucleosome core particle (rNCP) with a 151 base pair 601 Widom DNA as previously described (50,51). Briefly, the histones were purified from inclusion bodies under denaturing conditions on a 5 mL HiTrap SP FF (GE Healthcare) cation exchange column on a Next Generation Chromatograph (NGC, Bio-Rad). Fractions containing the purified histone were pooled and dialyzed 3 times into 4 L of water and 2 mM β-mercaptoethanol before lyophilization. The four histones were then unfolded into 20 mM Tris pH 7.5, 7 M Guanidine-HCl and 10 mM DTT and mixed in equimolar ratios prior to octamer refolding into 2 M NaCl, 10 mM Tris pH 7.5, 1 mM EDTA. Octamers were then purified on a Superdex 200 HiLoad 16/600 size exclusion column (GE Healthcare) and wrapped with the 151 base pair 601 Widom DNA to obtain rNCPs. Native gel analysis was used to validate the quality of the reconstitution.

### Histone labeling

The installation of a monomethyl-lysine analog at the mutated cysteine of the H3K_C_79 (C110A) histone was done as described (52) using the 2-chloro-N-methylethanamine hydrochloride (Toronto Research Chemicals C428323) to generate H3K_C_79^me1^ (C110A). The installation of the analog was confirmed by electrospray ionization mass spectrometry (ESI-MS) on a LC-ESI-QTOF Agilent 6538 mass spectrometer and immunoblotting against H3K79^me1^ (**Figure S2**).

## Supplementary Material Legends

**Figure S1:**
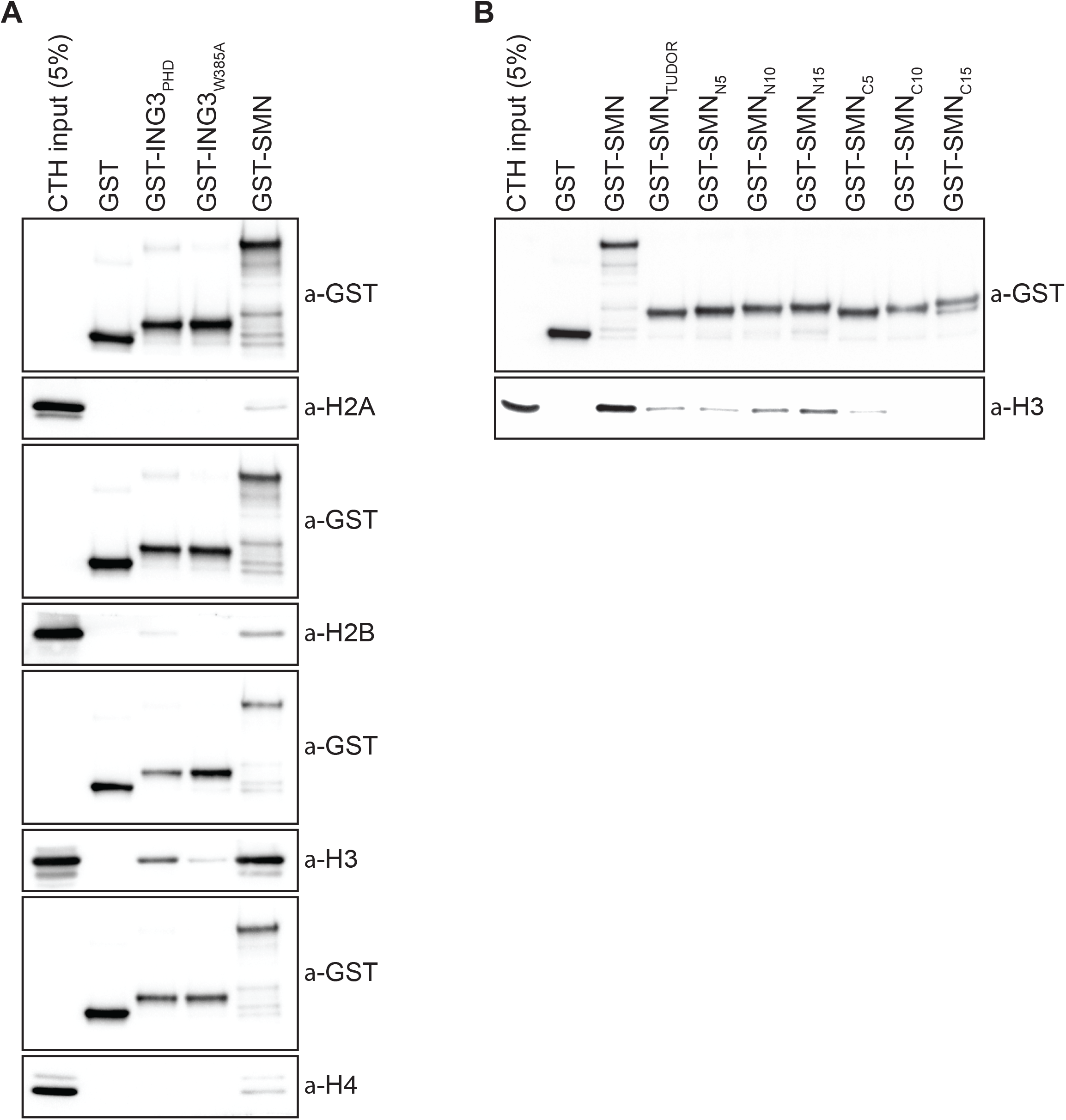
SMN associates with histone H3. (**A**) Samples from **Figure 1B**, but pulled down GST levels are shown for each immunoblots. (**B**) GST alone or GST-tagged indicated amino and carboxy terminal extensions on recombinant SMN_TUDOR_ were used in GST-pulldown assays in the presence of calf thymus histones (CTH). Pulldowns were analyzed by immunoblotting using the indicated antibodies.

**Figure S2:**
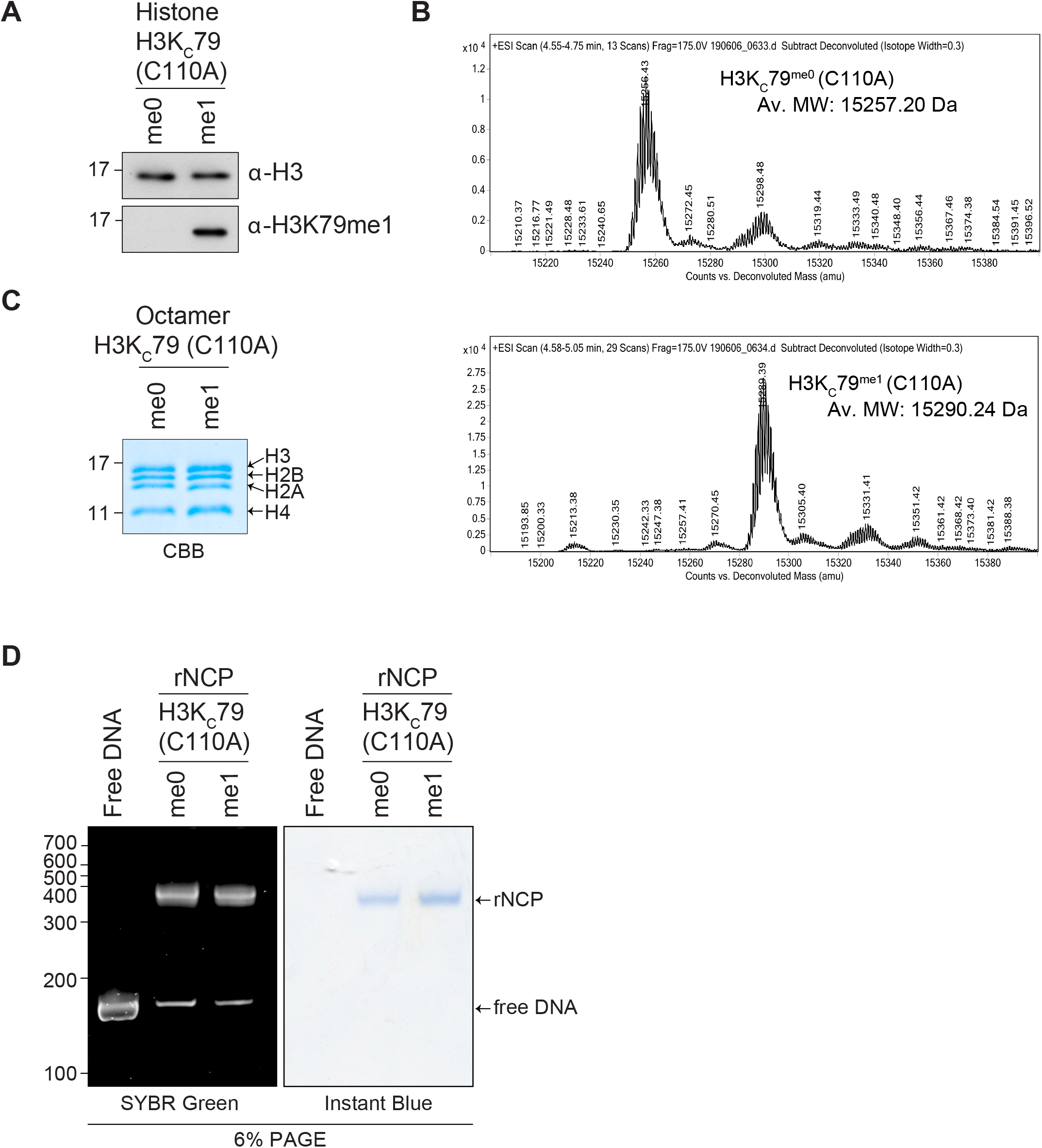
Analysis of rNCPs reconstitution with H3/K_C_29^me0^ (C110A) or H3/K_C_29^me1^ (C110A). Histone H3 H3K_C_79 (C110A) either unmodified or chemically modified with a monomethyl lysine analog, were analyzed by (**A**) immunoblotting using α-H3K79^me1^ or α-H3 and by (**B**) electrospray ionization mass spectrometry (ESI-MS) analysis. Total mass shift due to K_C_79 monomethylation: calculated +37; observed +33. (**C**) Octamers refolded the indicated histone H3 were separated on a 15 % SDS-PAGE and stained with Coomassie Brilliant Blue (CBB). (**D**) Reconstituted rNCP were run on native 6% retardation gel and sequentially stained by SYBR Green and InstantBlueTM (ISB1L, Sigma). Free 151 bp 601 DNA is used as a control.

**Table S1:** Summary of SMN association with various forms of histone H3. Qualitative estimates of the association between SMN and histone H3 (from CTH; Figure 4B), H3 monomethylated on K79 (from CTH; Figure 5F), and rNCP-H3K_C_79^me1^ (Figure 6).

| SMA mutants | Total H3 | H3K79<sup>me1</sup> | rNPC-H3K<sub>C</sub>79<sup>me1</sup> |
| --- | --- | --- | --- |
| W92S | - | - | - |
| V94G | + | + | + |
| G95R | - | + | - |
| Y109C | + | - | - |
| A111G | - | - | - |
| I116F | - | - | - |
| Y130C | + | + | - |
| E134K | + | + | + |
| Q136E | + | + | + |

## Acknowledgements

We thank Dr Faouzi Baklouti (DR2 Inserm) for proofreading and helpful discussions. This work was supported by AFM Téléthon grants awarded to PL (Plans stratégiques MyoNeurALP) as well as funding from the Joint Collaborative Research Program of University of Ottawa Centre for Neuromuscular Disease and Claude Bernard Université Lyon 1 Institute NeuroMyoGene awarded to OB and PL. JC is funded by a Canadian Institutes of Health Research (CIHR) grant (MOP 123381) and CureSMA Canada. AFT is funded by the Natural Sciences and Engineering Research Council of Canada (NSERC) Grant (RGPIN-2016-05844). AFT is a tier 2 Canada Research Chair in Molecular Virology and Genomic Instability and is supported by the Foundation J.-Louis Lévesque. MG is supported by a doctoral fellowship from the Fonds de Recherche du Québec - Santé (FRQS). PL is a CNRS research director.

